# Iron-dependent essential genes in *Salmonella* Typhimurium

**DOI:** 10.1101/159921

**Authors:** Sardar Karash, Young Min Kwon

**Affiliations:** Cell and Molecular Biology Program, University of Arkansas, Fayetteville, Arkansas; Department of Biology, College of Education, Salahaddin University, Erbil, Kurdistan, Iraq; Department of Poultry Science, University of Arkansas, Fayetteville, AR 72701

**Author notes:** Corresponding author: (YK).

**Keywords:** Essential genes, Iron-dependent essential genes, ROS, Tn-seq, Antibiotic targets, *Salmonella* Typhimurium

## Abstract

**Background:** The molecular mechanisms underlying bacterial cell death due to stresses or bactericidal antibiotics are complex and remain puzzling. Previously, it was shown that iron is required for effective killing of bacterial cells by numerous bactericidal antibiotics. Here, our high-resolution Tn-seq analysis demonstrated that transposon mutants of *S.* Typhimurium with insertions in essential genes escaped immediate killing or growth inhibition under iron-restricted conditions for approximately one-third of all essential genes.

**Results:** We grouped essential genes into two categories, iron-dependent and iron-independent essential genes. The iron-dependency of the iron-dependent essential genes was further validated by the fact that the relative abundance of these essential gene mutants increased further with more severe iron restrictions. Our unexpected observation can be explained well by the recently proposed common killing mechanisms of bactericidal antibiotics via production of reactive oxygen species (ROS). In this model iron restriction would inhibit production of ROS, leading to reduced killing activity following blocking of an essential function. Interestingly, the targets of most antibiotics currently in use clinically, whether bacteriostatic or bactericidal, are iron-dependent essential genes.

**Conclusions:** Our result suggests that targeting iron-independent essential genes may be a better strategy for future antibiotic development, because blocking these genes would lead to immediate cell death regardless of iron concentration. On the contrary, blocking iron-dependent pathways under iron limited *in vivo* environment could lead to reduced killing action, which might increase drug-resistance by mutagenic action of sublethal concentrations of ROS. This work expands our knowledge on the role of iron to a broader range of essential pathways, and provides novel insights for development of more effective antibiotics.

## Background

Essential genes encode the proteins that are essentially required for cell viability or growth. These genes have been exploited as pivotal targets for antibacterial drugs, because blocking their proteins cause cell impairment and ultimately growth inhibition or death of bacterial cells. Thus, nearly all antibiotics in clinical use target these essential pathways. However, for many natural antibiotics, the molecular targets remain unknown [1] and even if the target is known, in case of bactericidal antibiotics, the cellular events that follow in response to disruption of essential pathways leading to bacterial cell death remain puzzling.

Numerous studies have shown the role of reactive oxygen species (ROS) in cell death for eukaryotes as well as prokaryotes. In eukaryotes, apoptosis and necroptosis are associated with ROS [2, 3]. Ferroptosis is an iron-dependent nonapoptotic form of oxidative cell death in mammalian cancer cells. These cells die as a result of ROS accumulation and the death can be prevented via iron chelators [4]. In bacteria, contribution of ROS to cell death due to bactericidal antibiotics is supported by numerous studies. Kohanski et al. [5] proposed that bactericidal antibiotics, regardless of their molecular targets, induce production of ROS, which consequently damages biomolecules contributing to cell death, and demonstrated the death process can be mitigated via iron chelators. This model asserts that upon antibiotic-target interactions, consecutive specific intracellular events induce ROS formation, specifically hydroxyl radical, via Fenton reaction through the process that involves TCA cycle-NADH depletion and destabilization of Fe-S clusters [5, 6]. Furthermore, it was also shown that ROS generation elevates in bacterial cells by the attack of competitor bacteria or P1vir phage via type VI secretion system [7]. In addition, mammalian peptidoglycan recognition protein-induced bacterial killing requires ROS and the lethality of this protein can be inhibited via an iron chelator [8]. Immune cells also produce ROS to kill bacterial pathogens [9]. However, despite these numerous evidences on the role of ROS in bacterial cell death, it is unknown if this role of ROS can be generalized to all death process in bacterial cells, and if not, what the scope of the ROS-mediated cellular death process is.

A pathogenic bacterium can possess a few hundred essential genes that are indispensable for maintaining cell viability. Empirically, essential genes are defined by the genes that when inactivated lead to loss of cell viability. In *E*. *coli* Keio collection, single-gene deletions were made for all known open reading frames, excluding 302 genes which could not tolerate disruptions and these 302 genes were considered essential [10, 11]. On the other hand, random genome-wide transposon mutagenesis coupled with next generation sequencing (Tn-seq) is a powerful method to identify essential genes [12]. Tn-seq experiments have shown that there are 353 essential genes in *Salmonella* Typhimurium SL326 [13]; 461 in *Mycobacterium tuberculosis* H37Rv [14]; and 227 in *Streptococcus pyogenes* [15]. Recently, a team chemically synthesized *Mycoplasma mycoid* JCVI-syn3.0 based on 473 essential genes [16]. Clustered Regularly Interspaced Short Palindromic Repeats Interference (CRISPRi) was employed for phenotypic analysis of 289 essential genes in *Bacillus subtilis* that were identified by Tn-seq, and confirmed that approximately 94% of the putative essential genes were genuine essential genes [17].

Nearly all studies on defining essential genomes in bacteria have been conducted using stress-free nutrient-rich media for the given bacterial species under the assumption that a minimum set of the core essential genes would be best revealed under such “optimal” growth conditions. In this study, on the contrary, we analyzed our Tn-seq data to determine essential genes in *S.* Typhimurium under the stress conditions created by restricting iron concentrations using iron chelator 2,2’-Dipyridyl (Dip) ranging from 0 to 400 μM. Our initial focus was to identify conditionally essential genes required for fitness under iron-restriction conditions. However, we unexpectedly found that a considerable portion of the genes that are categorized as essential genes in LB media (no Dip) were rendered non-essential under iron-restriction conditions. Furthermore, the relative abundance of the transposon mutants in those essential genes increased with the increasing severity of iron restrictions. We reason that this finding has significant implications for the current efforts to overcome the crisis in public health due to increasing antibiotic resistance, and may provide valuable insights for future direction for the development of new antibiotics with inherently reduced chance of developing drug resistance. Therefore, in this study we focus on the analysis of the essential genes under iron-restricted conditions, and discuss the implications of our discovery.

## Results and Discussion

### Library selection and initial Tn-seq analysis

We constructed two genome-saturating Tn5 transposon libraries (Library–A and –AB) in which 92.6% of all ORFs had insertions (Table 1). To track the relative abundance of mutants in the libraries in response to iron restriction, appropriate library was inoculated into LB media supplemented with iron chelator 2,2’-Dipyridyl (Dip) at final concentrations of 0 (controls: LB-I, LB-II and LB-III), 100 (Dip100), 150 (Dip150), 250 (Dip250-I and Dip250-II), or 400 µM (Dip400) as shown in Fig. 1. The cultures were grown until the bacteria reached mid-log phase. Since the growth rate of *S.* Typhimurium was reduced by Dip treatment in a concentration-dependent manner (Fig. 2), the cultures were incubated for different lengths of time to reach the mid-log phase (Table 1). We also included Library-A itself (LB-I) for Tn-seq analysis as an additional control group (Fig. 1). From Library-AB, we obtained 273 million (M) sequence reads from Tn5 genomic junctions in the chromosome of *S*. Typhimurium for all conditions combined, and 185 M sequence reads were mapped to the completed genome of *S*. Typhimurium 14028 (Table 2).

**Fig.1.**
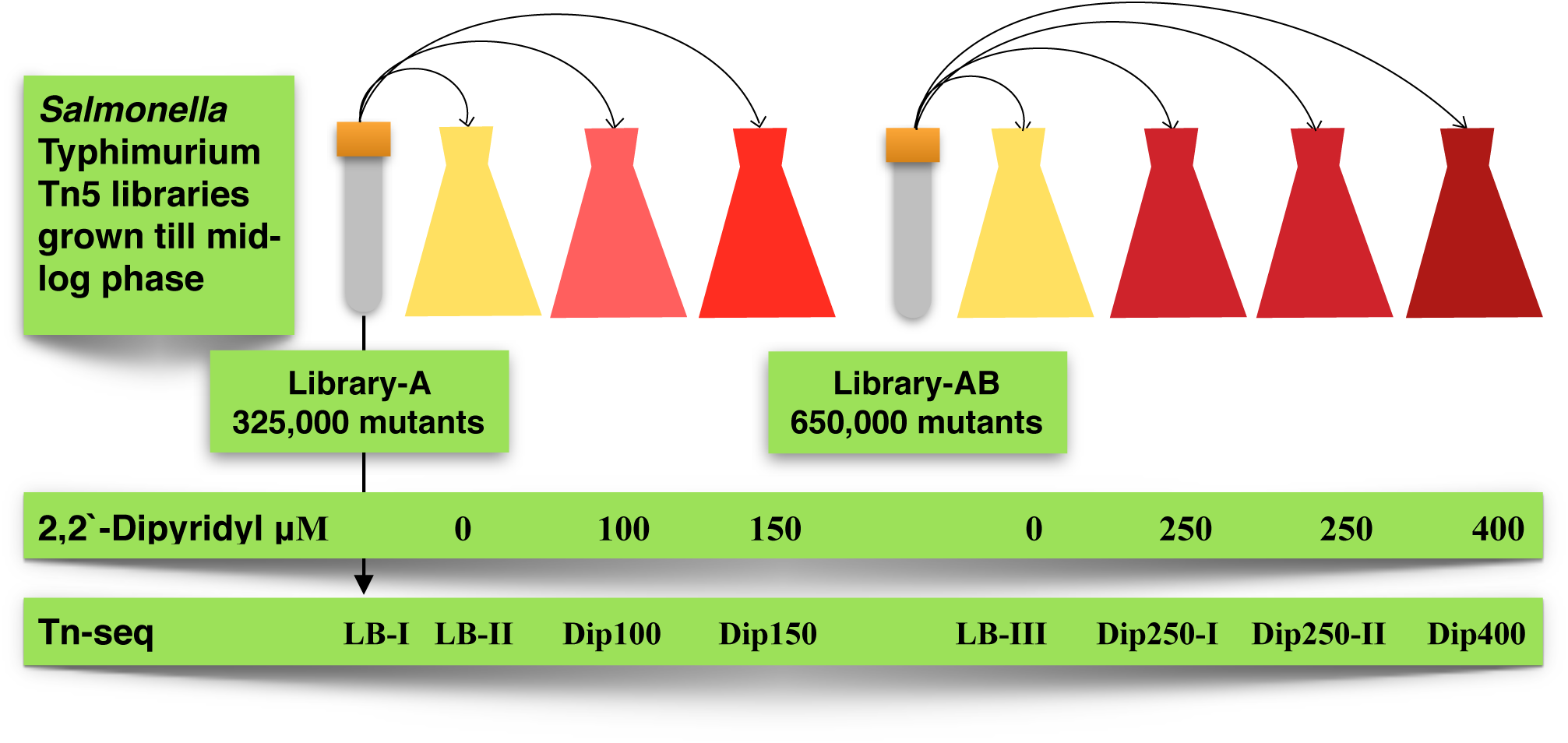
Schematic representation of study design. Transposon library-A inoculated to LB broth (LB-II) or the LB contained either 100 μM iron chelator Dip (Dip100) or 150 μM Dip (Dip150). LB-I was library-A that subjected to Tn-seq without growth. Transposon library-B inoculated to LB broth (LB-III) or the LB contained either 250 μM Dip (Dip250) or 400 μM Dip (Dip400). The cultures were grown till mid-log phase and then subjected to Tn-seq.

**Fig. 2.**
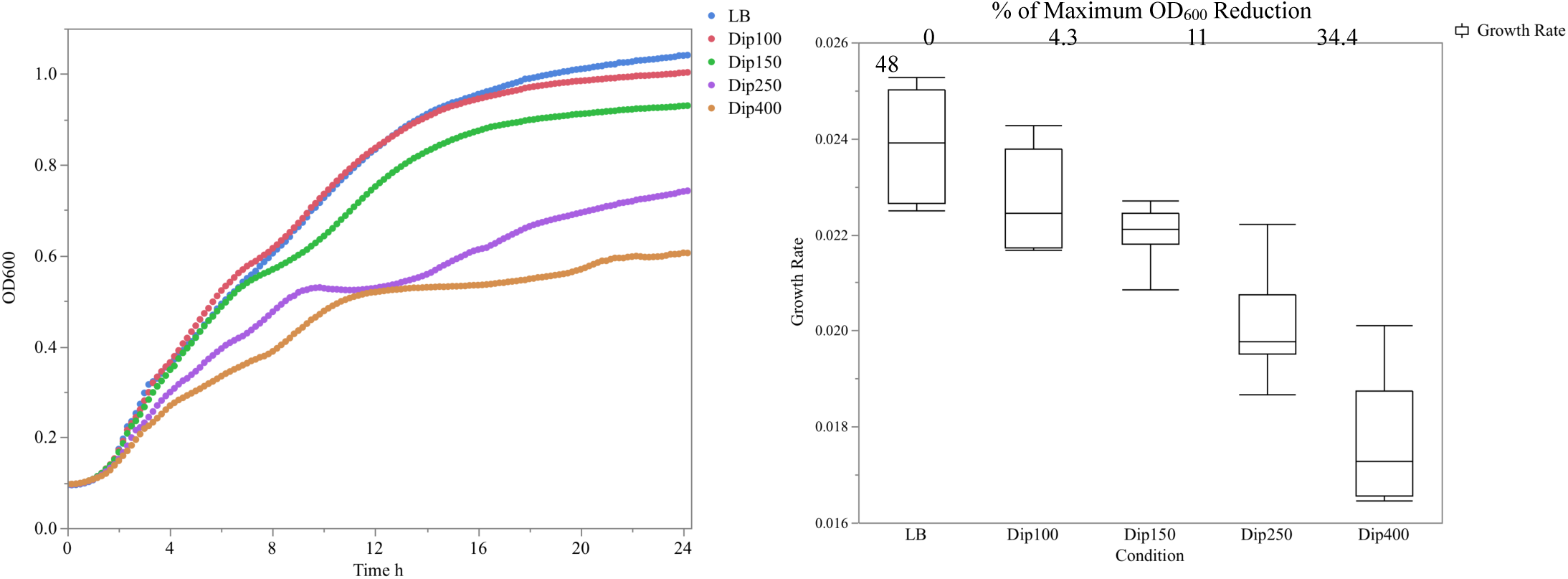
Effect of 2,2’-Dipyridyl (Dip) on *S.* Typhimurium growth. An overnight culture diluted 1:200 in LB broth supplemented with 100 μM Dip (Dip150), 150 μM Dip (Dip150), 250 μM Dip (Dip250), 400 μM Dip (Dip400), or Dip free (LB). The cultures were added to a 96-well plate and directly incubated at 37°C in a plate reader, reading OD600 every 10 minuets. The maximum OD600 reduction is shown as a percentage.

**Table 1.** Transposon inoculum densities and CFUs at mid-log phase. The seeding CFUs of all cultures counted following inoculation at time zero and at the mid-log phase when the growth stopped. OD600 measurements were used to monitor the growth. LB is broth free of Dip. Dip is abbreviation of iron chelator 2,2’-Dipyridyl in μM. The number with Dip is concentration of Dip.

| Table 1. Transposon inoculum densities and CFUs at mid-log phase. |  |  |  |  |
| --- | --- | --- | --- | --- |
|  | Seeding<br>CFUs/ml | CFUs/ml at mid-log phase | Time to reach mid-log phase h:min | OD<sub>600</sub> |
| **Library-A** | **325,000 mutant colonies recovered on 50 LB agar plates** |  |  |  |
| **LB-II** | 3,500,000 | 117,000,000 | 5 | 2.63 |
| **Dip100** | 3,500,000 | 190,000,000 | 5:35 | 2.61 |
| **Dip150** | 3,500,000 | 164,000,000 | 6:05 | 2.565 |
| **Library-AB** | **325,000 mutant colonies recovered on 50 LB agar plates + Library-A (total: 650,000 mutants)** |  |  |  |
| **LB-III** | 8,000,000 | 700,000,000 | 5:30 | 2.567 |
| **Dip250-I** | 8,000,000 | 2,000,000,000 | 10 | 2.446 |
| **Dip250-II** | 8,000,000 | 2,010,000,000 | 10 | 2.462 |
| **Dip400** | 8,000,000 | 550,000,000 | 24 | 1.843 |

**Table 2.** Sequence read counts used in this study. Total reads with Tn5 represent the sequence reads that passed the quality control and had sequence of Tn5. Extracted reads > 20 bp represent sequence reads that had trimmed C-tail (if present) and their length were above 20 nucleotides. Number of mapped reads, unique insertions in chromosome with mean length of mapped reads are shown. LB is broth free of Dip. Dip is abbreviation of iron chelator 2,2’-Dipyridyl in μM. The number with Dip is concentration of Dip.

| **Table 2. Sequence read counts used in this study.** |  |  |  |  |  |
| --- | --- | --- | --- | --- | --- |
|  | **Total reads**<br>**with Tn5** | **Extracted reads**<br>**>20bp** | **Mapped**<br>**Reads** | **Unique**<br>**Insertions** | **Mean genomic length**<br>**bp** |
| **Library-A** | **325,000 mutant colonies recovered on 50 LB agar plates** |  |  |  |  |
| **LB-I** | 18,225,644 | 14,437,819 | 12,289,451 | 115,784 | 93.8 |
| **LB-II** | 38,808,640 | 31,728,005 | 25,223,444 | 125,449 | 92.9 |
| **Dip100** | 25,788,698 | 21,034,947 | 16,991,894 | 117,474 | 93 |
| **Dip150** | 36,677,408 | 29,905,496 | 24,364,738 | 121,132 | 93.1 |
| **Library-AB** | **325,000 mutant colonies recovered on 50 LB agar plates + Library-A (total: 650,000)** |  |  |  |  |
| **LB-III** | 57,779,778 | 47,575,248 | 39,248,662 | 193,728 | 90.6 |
| **Dip250-I** | 29,832,849 | 25,082,465 | 21,096,630 | 179,562 | 90.1 |
| **Dip250-II** | 35,439,669 | 28,119,351 | 23,104,233 | 181,534 | 89.1 |
| **Dip400** | 30,028,187 | 26,382,625 | 23,135,546 | 169,666 | 91 |
| **Total** | 272,580,873 | 224,265,956 | 185,454,598 | 1,204,329 | 91.7 |

The accurate genome-mapping based on long Tn5 genomic junctions and high number of read counts allowed us to define essentiality and conditional essentiality of the genes with a high precision. Our initial goal in this study was to elucidate the conditionally essential genes that are required for fitness under different levels of iron restriction using Tn-seq. During the data analysis, however, we found the read counts corresponding to the mutants in numerous known essential genes increased significantly and consistently with iron restriction. This observation was in contrary to the currently accepted working definition of essential genes that cannot tolerate disruptions. It required further detailed analysis before we could accept this interesting, yet unexpected finding. Therefore, we have conducted a systematic analysis for essential genes and comparatively analyzed the results between iron-replete and iron-restricted conditions.

### Essential genome of *S*. Typhimurium in iron-replete and iron-restricted conditions

We used a rigorous analytical pipeline for essential gene identification as outlined in additional file 1. As a result, we identified 336 essential genes that are required for aerobic growth of *S*. Typhimurium 14028 in LB broths and on LB agar plates (Additional file 2). We compared the essential genes in *S*. Typhimurium 14028 to those in *S*. Typhimurium SL3261 identified by TraDIS [13]. Interestingly, out of 336 genes in our essential gene list, 265 (80%) orthologous genes in *S*. Typhimurium 14028 were also essential in *S.* Typhimurium SL3261 (Additional file 3). This is a significant overlap considering variations in the genetic backgrounds of the two strains. Further, KEGG pathway analysis recognized 306 out of the 336 genes, which were categorized into 23 essential pathways (Additional file 4).

We also used the same method to identify essential genes from the Tn-seq data obtained from iron-restricted conditions. Surprisingly, the number of essential genes under iron-restricted conditions decreased to 215 genes, which indicated that 121 genes (36%) of the 336 essential genes were considered non-essential under iron-restriction conditions (Additional file 5). The number of reads corresponding to the insertions in these 121 genes significantly increased under iron-restricted conditions: the average read counts in the 121 genes were 4.3 in LB-III whereas this elevated to 68 in Dip400 (Additional file 6). This is a clear evidence that the mutants of the 121 genes escaped immediate killing or growth arrest, and were able to multiply slowly under iron-restricted conditions. In another word, chelation of iron in the media allowed the mutants of these 121 genes to become no longer “essential” according to the working definition of an essential gene (Fig. 3, Additional file 7).

**Fig. 3.**
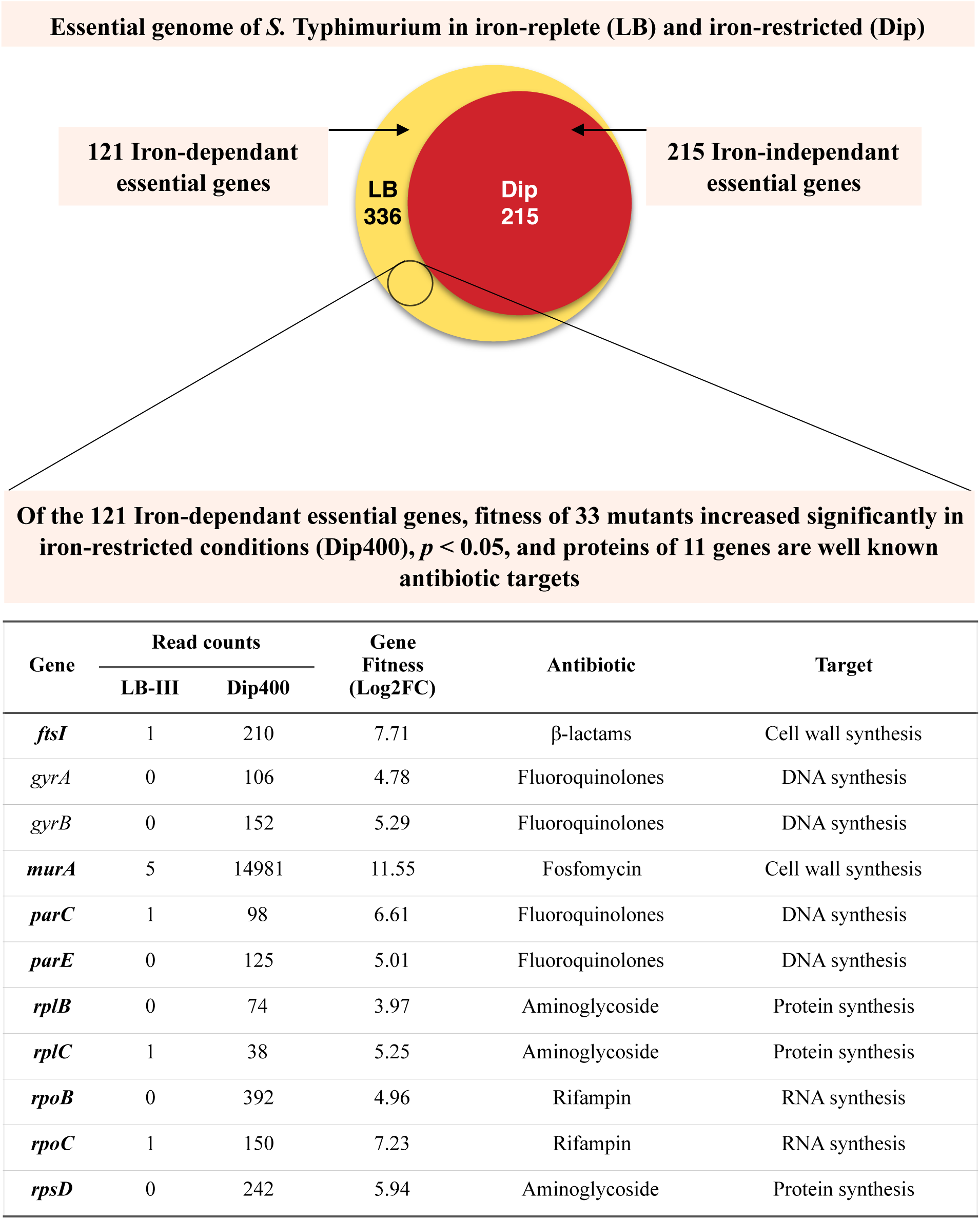
Iron-independent essential genes and iron-dependent essential genes of *Salmonella* Typhimurium. Essential genes of *S.* Typhimurium were identified using Tn-seq method in LB media (iron-replete) and LB media supplemented with 2,2’-Dipyridyl (Dip; iron-restricted). Among all 336 essential genes identified, 215 genes were essential under both conditions (iron-independent essential genes), while 121 genes were essential only in iron-replete condition (iron-dependent essential genes). The iron-dependent essential genes included 11 genes encoding well known antibiotic targets as shown in the table below.

### Fitness of the mutants of iron-dependent essential genes

We next measured the changes of mutant fitness for the 121 genes in the presence of high concentrations of Dip using Dip250-I, Dip250-II and Dip400 Tn-seq data as outputs in comparison to LB-III as the input. Strikingly, the mutant fitness of 97 out of 121 genes (78%) increased numerically (fitness expressed in Log_2_ fold change in read counts) in Dip400, and 8 other genes showed increased mutant fitness in Dip250-I, Dip250-II or both. Further comparative analysis revealed that 33 essential genes showed significant increase in fitness in the presence of Dip, including *gyrA*, *gyrB*, and *ileS* (*p* value < 0.05; Fig. 4). For these 33 genes, the mutants showed at least Log_2_ fold change in read counts ≥ 3.64 in Dip400 in comparison to LB-III. Interestingly, *gyrA*, *gyrB* and *ileS* were not in the list of 121 iron-dependent essential genes (e.g. considered as iron-independent essential genes), whereas fitness of these mutants increased significantly by iron restriction. In the case of *murA*, the read counts increased from 5 (LB-III) to 14,981 (Dip400), exhibiting highest increase in abundance (Fig. 4). These are additional strong evidences supporting that the genes with positive fitness in iron-restricted conditions are rendered nonessential and multiplied in iron-restriction conditions.

**Fig. 4.**
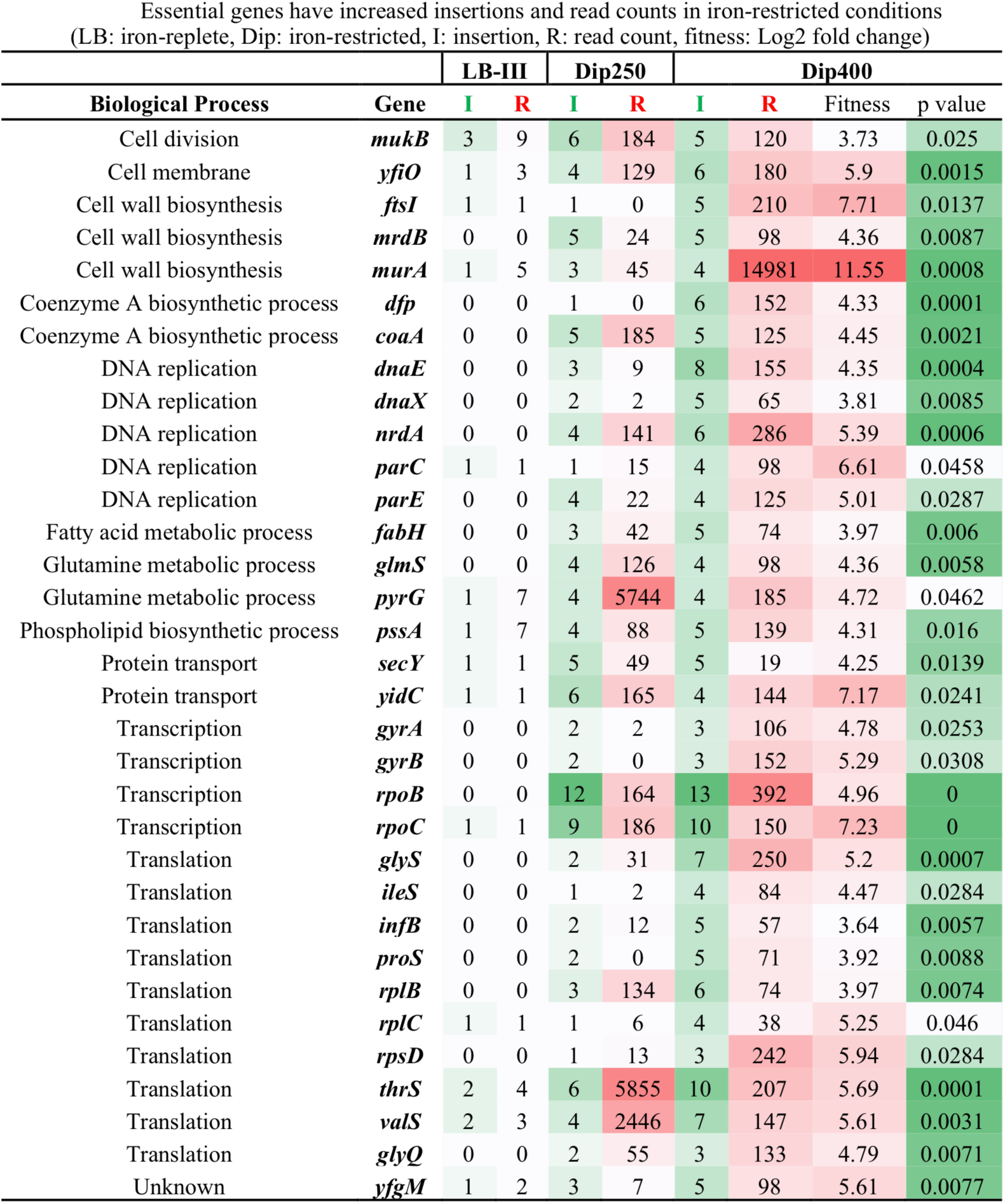
Iron-dependent essential genes have increased insertions and read counts in iron-restricted conditions. *Salmonella* Typhimurium transposon mutants were grown in LB media (iron-replete) and LB media supplemented with Dip (iron-restricted). Tn-seq analysis identified 121 genes including these genes as essential in iron-replete, but not in iron-restricted conditions. The fitness of these 33 genes increased in Dip400. Gene fitness is Log_2_ fold change of sequence reads in Dip400 in comparison to LB-III.

### Iron-dependent essential genes are not condition-specific essential genes

It is well known that the essential genes are operationally defined depending on the variations in the specific optimal growth conditions used for the experiment. We then asked if our discovery could be considered as a general extension of the concept to iron-restricted conditions. To answer this question, we closely examined our Tn-seq data that were generated for other stress conditions such as H_2_O_2_[18] and H_2_O_2_ coupled with Dip (unpublished). However, we could not find any similar patterns that a significant portion of the essential genes in LB medium became nonessential under stress conditions. We also examined other studies on identification of essential genes in other bacteria, including *Salmonella* Typhimurium SL326 [13], *Mycobacterium tuberculosis* H37Rv [14] and *Mycoplasma mycoid* JCVI-syn3.0 [16] as well as *E*. *coli* [10]. Nearly all orthologous essential genes in these bacteria, specifically the 33 genes, barely tolerated insertions and thus considered essential in their stress-free conditions (Additional file 8). Further, Lee et al [19] used Tn-seq to identify essential genes in *Pseudomonas aeruginosa* under 6 different conditions, and found that the 352 genes are general essential genes important in all 6 different media, while 199 essential genes were condition-specific. The condition-specific essential genes constitute 11-23% essential genes depending on the growth medium. It is important to note that all these 6 media support the growth of *P. aeruginosa* well, and thus are not considered as stress conditions. In contrast, in our study 36% of the essential genes in LB medium became nonessential under iron-restricted stress conditions, and there were no condition-specific essential genes for iron-restriction condition (Fig. 3). Therefore, we reason that these 121 genes should be still considered as general essential genes, instead of LB-specific essential genes. In other words, these 121 genes are general essential genes, but became non-essential under the iron-restricted condition due to the critical role iron plays in broad range of death processes in *S.* Typhimurium following inactivation of the essential functions.

### Validation of the Tn-seq results

We next asked whether the increase of insertions and read counts for the 121 essential genes under iron-restricted conditions were due to a bias in Tn-seq data analysis. We conducted the analysis for identification of essential genes in this study without data normalization. Typically normalization of read counts is critical for reliable identification of conditionally essential genes for the mutant libraries before and after the selection. On the contrary, essential gene analysis is usually conducted without normalization. It is because the process is based on one Tn-seq profile from the condition of interest under which the essential genes are studied, and the information on insertion sites is critical for discovery of essential genes, while relative abundance of mutants is not. In this study, there were differences in read counts in Tn-seq conditions (Table 2). Furthermore, when focused on the reads in ORFs only, the total read counts of LB-III was ∼30 millions (M) versus ∼16 M in Dip400, excluding intergenic regions (Additional file 9). Interestingly, the insertions in the two genes, *STM14_2422* (*umuC*) and *STM14_2428*, consumed 8.7% of all reads in LB-III and 27% in Dip400. Consequently, excluding the reads from these two genes, an insertion in LB-III is expected had on average 227 reads, while 100 reads for an insertion in Dip400 (Additional file 9). This indicates that the bias in read counts was in favor to not see insertions and read counts in essential genes in iron-restricted conditions (Dip400). Despite this bias in read counts, which was further intensified by reads from these two genes, the read counts in the 121 iron-dependent essential genes were higher in Dip400 compared to LB-III. This is a strong evidence that the mutants in these 121 genes did indeed multiply slowly in iron-restricted conditions.

### Iron-dependent and -independent essential genes

We came to the conclusion that iron-restriction alleviated the immediate killing or growth arrest of the mutants in the 121 iron-dependent essential genes, allowing growth of the mutants, although the underlying mechanism(s) is unknown. Here, we propose to divide the essential genes in *S.* Typhimurium for convenience into two categories according to the dependency of their gene essentiality on iron: iron-dependent and iron-independent essential genes (Fig. 3). The 121 essential genes that allowed growth of the mutants under iron-restricted conditions are iron-dependent essential genes (Fig. 3, Fig. 5, and Additional file 6). On the contrary, the 215 essential genes that did not tolerate growth of the mutants regardless of iron concentrations are iron-independent essential genes (Fig. 3, Fig. 5, and Additional file 5).

**Fig. 5.**
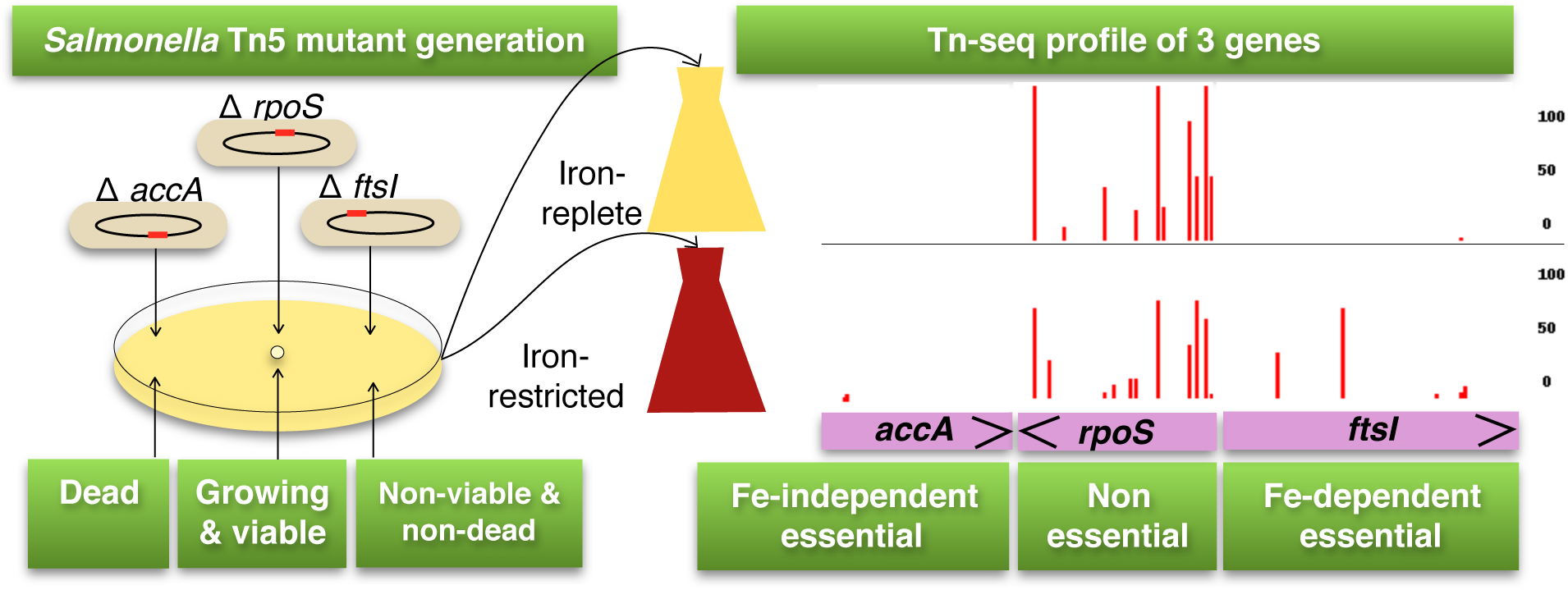
Identification of iron-dependent essential genes by Tn-seq. *Salmonella* Typhimurium Tn5 mutant library was inoculated to iron-replete media (LB media) and iron-restricted media (LB supplemented with an iron chelator 2,2’-Dipyridyl). The cultures were grown till mid-log phase and then Tn-seq was used to identify essential under each condition. The essential genes that were considered essential only in iron-replete condition are iron-dependent essential genes.

One note for caution for this classification is that for any iron-independent essential genes reported in this study the classification can be temporary. It is because the decision on gene essentiality can be dependent on the sequencing depth, and a substantial increase in the sequencing depth may reclassify some of the iron-independent genes as iron-dependent essential genes.

Then, we compared the average read counts between iron-independent and iron-dependent essential genes under iron-replete or iron-restricted condition. For the 215 iron-independent essential genes, their average read counts were 9.6 in Dip400 (Additional file 5). On the contrary, for the 121 iron-dependent essential genes, their average read counts were 67.9 in Dip400 (Additional file 6). Whereas in iron-replete condition, LB-III, the average read counts were only 2.2 for the 215 iron-independent essential genes and 4.3 for 121 iron-dependent essential genes. In addition, we asked if the 121 iron-dependent essential genes represent unique pathways in comparison to iron-independent essential genes. The result of KEGG pathway analysis showed that iron-independent and dependent genes are represented in 15 and 10 pathways, respectively, with 7 overlapping ones (Fig. 6).

**Fig 6.**
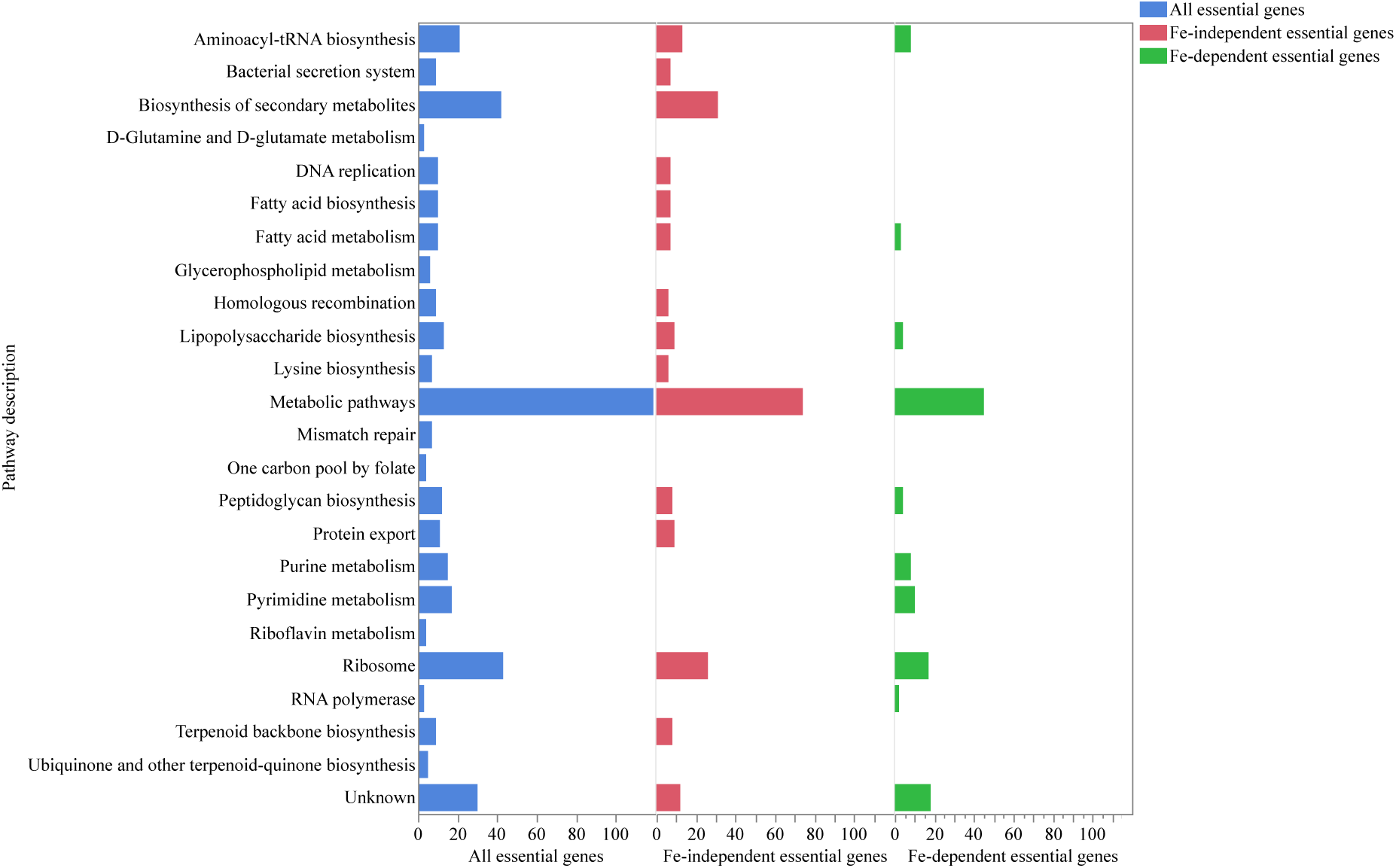
Fe-dependent essential pathways in *Salmonella* Typhimurium identified by Tn-seq. Transposon libraries were grown in iron-replete and iron restricted conditions. Essential genes identified for both conditions. The genes that were essential in iron-replete condition but not in iron-restricted conditions were considered Fe-dependent. KEGG pathway analysis was used for pathway description.

**Fig. 7.**
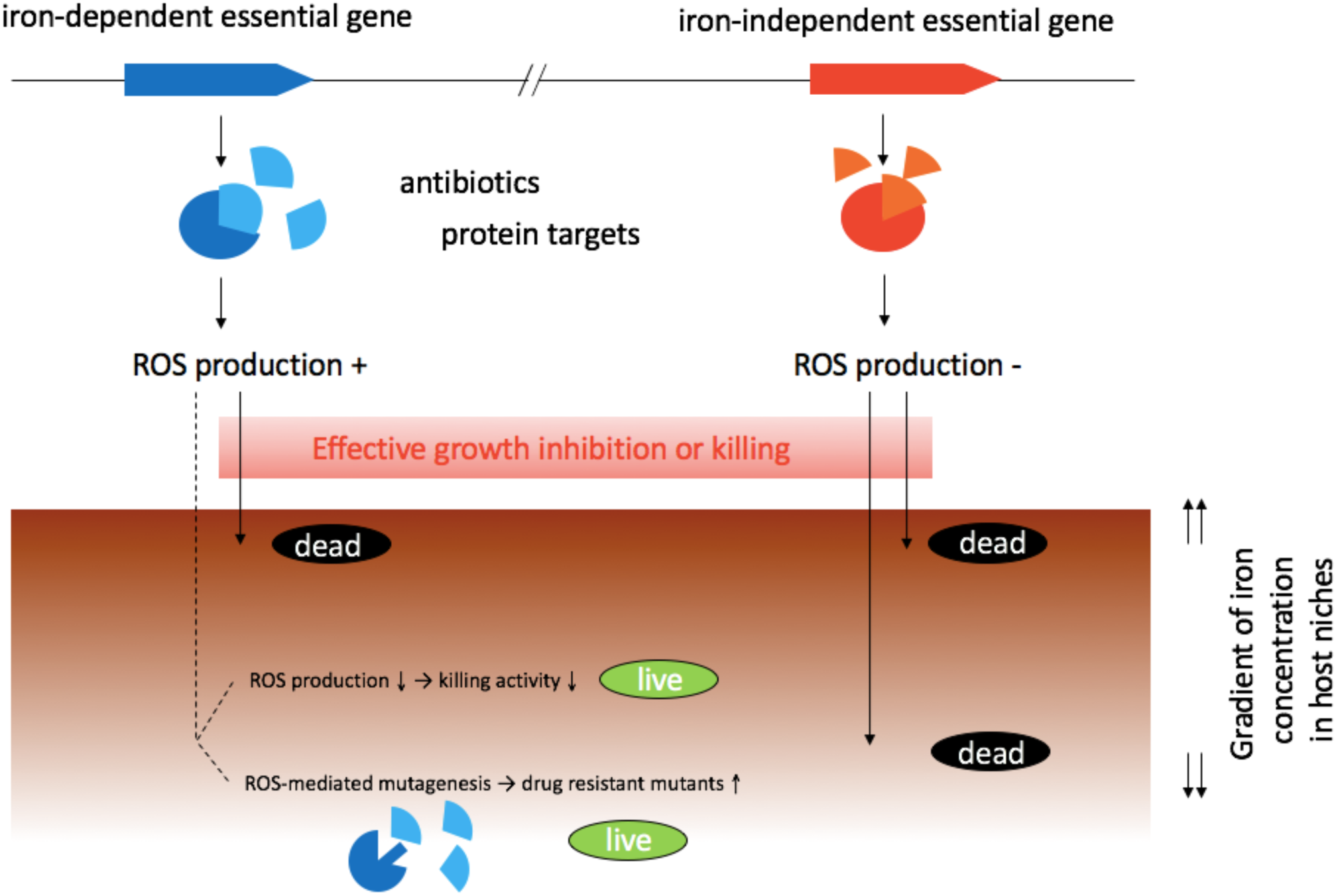
Iron-independent essential genes may serve as better targets for development of new antibiotics. The proposed model indicates that targeting iron-independent essential pathways would lead to effective killing or growth inhibition of *Salmonella* Typhimurium even in iron-restricted host niches.

It is also important to note that all known targets of antibiotics, with only one exception of colistin, are encoded by iron-dependent essential genes (Fig. 3). It is uncertain if this strong bias toward iron-dependent genes reflects the preference of the antibiotic-producing microorganisms in nature to target these essential genes. If that’s the case, it would be very intriguing to explore the possible mechanistic reasons for the bias in relation to the roles of antibiotic production in the natural environments.

### Fitting iron-dependent essential genes in ROS-mediated antibiotic killing model

It has been well established that iron contributes to cell death due to various causes. 2,2’-Dipyridyl iron chelator inhibits the Fenton reaction, which is required for ROS generation [5, 20]. We speculate that following disruption of the essential genes with transposon insertions, ROS might contribute to the death or growth arrest depending on the target genes, in addition to the disruption of the essential protein functions. We reason that our results have significant implications in understanding and expanding the current model of ROS-mediated common killing mechanisms of bactericidal antibiotics.

Since its first proposal by Kohanski et al. [5], this hypothesis has been substantiated by numerous studies using different bacterial species and bactericidal antibiotics [21–24]. Traditionally, the mechanisms of antibiotic action have been studied largely in terms of antibiotic-target interactions. However, numerous researches supporting the ROS-mediated killing mechanism have shown that the interaction of antibiotic-target leads to production of ROS, contributing to the killing activity mediated by direct blocking of the essential pathways for living cells. In our study, we did not use bactericidal antibiotics to disrupt an essential pathway. However, inactivation of the essential gene functions was achieved in a permanent and irreversible manner through Tn5 insertions in the essential genes, instead of reversible antibiotics treatment. In most studies focused on understanding the killing mechanisms of bactericidal antibiotics, the experiments were conducted using optimized sub-lethal concentrations of relevant antibiotics to allow reliable measurements of quantitative changes in killing effects (e.g. by addition of iron chelator) [5]. However, inactivation of the essential protein functions by Tn5 insertions in our study is expected to exhibit the lethal effect equivalent to or even stronger than that caused by high lethal concentrations of bactericidal antibiotics. This situation may limit the effect of iron chelation to very small, making it difficult to detect and measure reduced killing by iron chelator. However, the Tn-seq procedure conducted with deep sequencing in combination with the high concentrations of Dip (up to 400 μM, which is much higher than typically used) employed in this study allowed to detect the small reductions in killing or growth arrest under the iron-restricted conditions.

Until now the ROS-mediated killing mechanism has been studied and discussed in the context of a few bactericidal antibiotics and their molecular targets. Our Tn-seq data show that this ROS-mediated killing mechanism might be linked at least to one-third of the essential genes in *S.* Typhimurium, thereby potentially expanding the ROS-mediated lethal pathway as a more general mechanism connected to a broad range of basic essential pathways in *S.* Typhimurium.

### Iron-independent essential genes may be better targets for antibiotic development

We believe that our findings have profound implications for understanding current crisis in public health due to the rapid increase in antibiotic resistance to most antibiotics in clinical use. Our deeper understanding of the underlying mechanisms would help in development of novel antibiotics with inherently reduced chance of developing drug resistance. Here, we propose that iron-independent essential genes may serve as better targets for antibiotic development for the following reasons: it has been shown that there are two opposing aspects of ROS-mediated killing mechanism. When ROS production is high, it would lead to facilitated killing of bacterial cells. On the contrary, when ROS production is low, it would lead to production of resistant mutants through mutagenic action of ROS on DNA [25]. When *Salmonella* infects the host, iron-restricted host niches would suppress ROS production from bactericidal antibiotics, which would in turn reduce ROS-mediated killing process and thus overall killing effect by the antibiotics. Furthermore, depending on the iron restriction levels, and reduced local concentrations of antibiotics, it might allow production of low levels of ROS, which might facilitate production of antibiotic resistant mutants through its mutagenic action on DNA. In contrast, if a certain antibiotic targets an iron-independent essential pathway (such as those encoded by the 215 iron-independent essential genes discovered in this study), we speculate that since ROS production is not a part of their lethal processes, blocking the pathways will lead to iron-independent killing, without increasing the chance of developing antibiotic resistant population via mutagenic action of ROS [25].

Our Tn-seq results show clearly that mutants of iron-dependent essential genes can grow slowly in iron-restricted conditions, and the same phenomenon may happen in the host tissues, because iron-restriction by host is a vital mechanism to combat the pathogen. As a result, it may be hard to completely kill, and eliminate *Salmonella* by targeting iron-dependent essential genes. Conversely, blocking iron-independent essential pathways would allow immediate killing of *Salmonella* regardless of iron concentration. Thus, the possibility will be higher to eradicate this pathogenic bacterium by targeting the iron-independent essential pathways in comparison to iron-dependent essential pathways as is the case for all antibiotics in clinical use except colistin.

A mechanism that bacteria exploit for antibiotic resistance is alteration of drug interaction site. Our results emphasize that the majority of genes encoding currently known drug targets are iron-dependent essential genes (Fig. 3). Prevalence of antibiotic resistance in clinical isolates due to mutations in drug targets has been rising. Mutations in a peptidoglycan synthesis gene *fts* which is target of β-lactams in *Haemophilus influenzae* cause resistance to antibiotics [26, 27]. *E. coli* strains harboring mutations in *murA* are resistant to fosfomycin [28]. UDP-N-acetylglucosamine enolpyruvyl transferase (MurA) catalyzes the reaction in the first step of biosynthesis of peptidoglycan in bacterial cell wall, and the protein is the target of fosfomycin [29]. Our Tn-seq results show that *murA* mutants did grow very well in iron-restricted conditions and the mutant had 14,981 read counts in Dip400 but there were only 5 reads of this mutant in LB-III (Fig. 4). It has been reported that *Pseudomonas putida* develops intrinsic fosfomycin resistance due to the presence of a salvage pathway that bypasses *de novo* biosynthesis of MurA [30]. Since *murA* is an iron-dependent essential gene in our study, we reason that almost all *murA* mutants died effectively in LB-III because of the contribution of ROS in the death process. However, in Dip400, reduced ROS formation and the salvage pathway biosynthesis of MurA might have caused *S*. Typhimurium to grow well. Further, Fluoroquinolone-resistant bacteria are also present in clinical isolates due to mutations in drug targets, *gyrA*, *gyrB*, *parC*, and *parE*, in pathogens such as *Shigella flexneri* [31], *Salmonella* Typhi [32], and group B *Streptococcus* [33]. Rifampin-resistant *Mycobacterium tuberculosis* isolates are associated with mutations in their targets, *rpoB* and *rpoC* [34, 35]. Mutations in *rplC* contributed to *Staphylococcus aureus* resistance to linezolid in a clinical isolate [36]. Finally, Telithromycin resistant-isolates of *S. aureus* due mutations in a ribosomal gene, *rplB*, were detected *in vitro* [37]. All together, these antibiotic targets, which are iron-dependent essential genes, can mutate and alter the structure of corresponding proteins in order to evade lethal interactions with the antibiotics.

One example of an antibiotic targeting a protein encoded by iron-independent gene is colistin. Colistin (polymyxin E) is a last resort antibiotic for treatment of infections caused by multidrug resistant Gram-negative bacteria [38]. This bactericidal drug interacts with the lipid A moiety of lipopolysaccharide (LPS) and ultimately causes membrane lysis [39]. We showed that the genes encoding the molecular targets of Colistin, *lpxABCDHK* are iron-independent essential genes. Over the last 60 years, Colistin has been used for fighting infectious diseases, but it has been used with caution and limitation due to its known toxicity. This resulted in less frequent use of Colistin, which is considered as the main reason why bacterial resistance is low for Colistin. Our Tn-seq results indicate that disruption of LPS is lethal in *S*. Typhimurium and there is no contribution of ROS to death process caused by Colistin. However, one study demonstrated that Colistin induced *Acinetobacter baumannii* killing through ROS production [40]. The contradicted findings are not surprising, considering that there has been continued debates on the common antibiotic killing mechanism via ROS. Although this model is widely accepted, a few studies challenged it [41, 42].

## Conclusion

In this work we employed Tn-seq to elucidate the essential genes exhibiting iron-dependent and iron-independent mutant phenotypes in *S.* Typhimurium. Our Tn-seq data indicated that when transposon mutant library was treated with an iron chelator, the mutants of approximately one-third of essential genes escaped immediate growth inhibition or killing, and multiplied slowly, increasing their relative abundance. Based on this observation, we speculate that the iron chelator reduces ROS formation via inhibition of the Fenton reaction, thereby lessening the lethal outcomes following inactivation of the essential functions. Accordingly, we divided 336 essential genes in *S.* Typhimurium 14028 into iron-dependent vs. iron-independent essential genes depending on their dependency of the essentiality on iron concentration. We propose that iron-independent essential genes may serve as better targets to develop new antibiotics, because targeting these pathways would lead to immediate killing or growth inhibition regardless of local iron concentrations in the host tissues. The proposed model illustrating our concept of iron-dependent and iron-independent essential genes is summarized in Fig 5. Obviously, further studies providing experimental evidences on the direct involvement of ROS production in the process would be necessary to support our model. Due to the difficulty associated with the phenotype of essentiality, the use of inducible knockouts such as CRISPRi [17] would be helpful in elucidating the role of ROS in the process following inactivation of the essential pathways. It would be also interesting to see if we could come to a similar conclusions using the same Tn-seq approach, when thiourea, hydroxyl radical scavenger, is used in place of Dip to suppress ROS production or the mutants are cultured anaerobically.

This study provides new insights on the previously unknown aspect of essential pathways in *Salmonella.* Although further studies are needed to gain better understanding on the scope of this observation (e.g. other bacteria), and to establish mechanistic basis of the iron-dependent essential genes, this study points us to a new direction of research that would be important to understand and overcome current crisis of antibiotic drug resistance.

## Methods

### Bacterial strains and Tn5 mutant library construction

*Salmonella* enterica subsp. enterica serovar Typhimurium ATCC 14028 with spontaneous mutation conferring resistance to nalidixic acid (NA) was used in this study. All procedures involving this pathogen (Biosafety level 2) were conducted according to the protocol approved by Institutional Biosafety Committee (IBC) at the University of Arkansas. Transposon mutant libraries were prepared as previously described by Karash et al., [18]. Briefly, *S.* Typhimurium ATCC 14028 was subjected to transposon mutagenesis by biparental mating using *Escherichia coli* SM10 λ*pir* carrying a pBAM1 transposon-delivery plasmid vector [43] as the donor. An equal volume of overnight growth cultures of the donor and recipient bacteria (*S.* Typhimurium) were washed with 10 mM MgSO_4_ and concentrated on the nitrocellulose filter, which was then incubated for 5 h at 37°C on a surface of LB agar plate. After the incubation, the cells were washed with 10 mM MgSO_4_ and plated on LB agar plates containing 50 µg/ml NA and 50 µg/ml Kanamycin (Km). The plates were incubated at 37°C for 24 h. Then, the colonies were scrapped off, added into LB broth supplemented with 7% DMSO, and stored at −80°C in aliquots. We constructed two mutant libraries, A and B. Each library contains approximately 325,000 mutants.

### Growth rate measurement for *S.* Typhimurium

A single colony of *S.* Typhimurium ATCC 14028 was inoculated into 2 ml LB broth medium in a 5 ml tube and incubated overnight (∼16 h). Freshly prepared LB broth media supplemented with different concentrations of 2,2’-Dipyridyl (Dip) were inoculated with the overnight culture at a 1:200 dilution. The cultures were immediately added into a 96-well microplate (200 μl/well) and incubated in a TECAN Infinite 200 microplate reader at 37°C (shaking amplitude of 1.5 mm, and shaking duration of 5 s), and OD_600_ was measured every 10 min for 24 h. The data were used to determine lag time phase, growth rate, and maximum OD_600_ for each concentration using GrowthRates script [44].

### Selection of mutant libraries for Tn-seq analysis

An aliquot of transposon libraries in stock was thawed at room temperature and diluted 1:10 in LB broth. The library was incubated at 37°C with shaking at 225 rpm for an hour and then washed twice in PBS. The library-A was inoculated to 20 ml LB broth (LB-II) and LB supplemented with either 100 (Dip100) or 150 μM Dip (Dip150) in a 300 ml flask, with seeding CFUs of 3.5 x 10^6^ per ml. We also included LB-I for which Library-A was directly subjected to Tn-seq after activation and washing. In addition, to accomplish a higher saturation level, Library-A was combined with Library-B (termed Library-AB; Fig. 1).

Library-AB was inoculated to 20 ml LB broth in a 300 ml flask (LB-III) and LB supplemented with either 250 (Dip250-I and Dip250-II) or 400 μM Dip (Dip400), at seeding CFUs of 8 x 10^6^ per ml as described above. Dip100, Dip150, Dip250-I, Dip250-II, and Dip400 along with LB-II and LB-III were incubated at 37°C with shaking at 225 rpm until the cultures reached mid-log phase (OD_600_ of ∼2.7). Then, the cultures were immediately centrifuged to get the cell pellets, which were stored at -20°C for downstream analysis.

### Preparation of Tn-seq libraries for HiSeq sequencing

The preparation of Tn-seq libraries was performed as previously described by Karash et al., [18]. Briefly, genomic DNA was extracted for each sample using DNeasy Blood & Tissue kit (Qiagen), and quantified using Qubit dsDNA RB Assay kit (Invitrogen). To remove the pseudo Tn5 mutants generated by chromosomal integration of pBAM1, genomic DNA was digested with PvuII-HF (New England Biolabs), and purified with DNA Clean & Concentrator-5 kit (Zymo Research). Then, a linear PCR extension was performed using Tn5-DPO (5’-AAGCTTGCATGCCTGCAGGTIIIIICTAGAGGATC-3’). The PCR reaction was performed in a 50 μl reaction containing GoTaq Colorless Master Mix (Promega), 20 μM Tn5-DPO primer, 100 ng gDNA, and H_2_O. The PCR cycles consisted of 95°C for 2 min, followed by 50 cycles at 95°C for 30 sec, 62°C for 45 sec, and 72°C for 10 sec. The PCR product was purified with the DNA Clean & Concentrator-5 kit. The C-tailing reaction was conducted with terminal transferase (TdT; New England Biolabs), CoCl_2_, dCTP, ddCTP, TdT buffer, and the purified linear PCR product. dCTP and ddCTP were included at the molar ratio of 20:1. The mixture was incubated at 37°C for 1 h, which was followed by 10 min incubation at 70°C. The C-tailed product was purified. Next, the exponential PCR was performed with P5-BRX-TN5-MEO primer, AATGATACGGCGACCACCGAGATCTACACTCTTTCCCTACACGACGCTCTTCCGATCTNNNNAG-BC-CCTAGGCGGCCTTAATTAAAGATGTGTATAAGAG (where “BC’ denotes different sample index barcodes of 8nt long) and P7-16G primer, CAAGCAGAAGACGGCATACGAGCTCTTCCGATCTGGGGGGGGGGGGGGGG. The PCR reaction was performed in a 50 μl reaction containing GoTaq Green Master Mix, P5-BRX-TN5-MEO primer, P7-16G primer, purified C-tailed Tn5-junction fragments, and H_2_O; the PCR cycles consisted of 95°C for 2 min, followed by 30 cycles of 95°C for 30 sec, 60°C for 30 sec, and 72°C for 20 sec, with the final extension at 72°C for 5 min. Then, the 50 μl PCR products were run on an agarose gel and the DNA fragments of size 325–625 bp were extracted using Zymoclean Gel DNA Recovery kit (Zymo Research). The DNA libraries were quantified using Qubit dsDNA RB Assay kit. The DNA libraries were combined and sequenced on a flow cell of HiSeq 3000 using single-end read and 151 cycles (Illumina) at the Center for Genome Research & Biocomputing in Oregon State University.

### Analysis of Tn-seq data

The HiSeq sequence results were downloaded onto High Performance Computing Center (AHPCC) at the University of Arkansas. The libraries were de-multiplexed using a custom Python script. The script searched for the unique 8-nucleotide barcode for each library for perfect matches. The transposon genomic junctions were extracted using Tn-Seq Pre-Processor (TPP) tool [45]. The TPP searched for the 19 nucleotide inverted repeat (IR) in a fixed sequence window and identified five nucleotides (GACAG) at the end of the IR sequence, allowing one nucleotide mismatch. The genomic junctions that start immediately after the GACAG were extracted and the C-tails were removed. The junction sequences of less than 20 nucleotides were removed and remaining junction sequences were mapped to the *Salmonella* enterica serovar Typhimurium 14028s genome and plasmid using BWA-0.7.12 [46]. The TPP counted and reported the number of total sequence reads after filtering, total mapped read, and total unique insertions in each library.

### Identification of essential genes

LB-I, LB-II, and LB-III were analyzed to identify the essential genes in *S*. Typhimurium 14028. We used two different tools for Tn-seq essential gene analysis. First, TRANSIT [45] analysis of essentiality on gaps in entire genome was conducted using tn5gaps algorithm. The 5% of N-terminal and 10% of C-terminal of open reading frames (ORF) were removed and even insertions with only one reads were considered for the analysis. The gene was considered essential if its *p* value ≤ 0.05. Secondly, Tn-Seq Explorer [47], was used for essential gene analysis by applying a 550 window size. The 5% of N-terminal and 10% of C-terminal ORFs were removed and insertions with only one reads were included in the analysis. The gene was considered essential if its Essentiality Index was ≤ 2. Then, the essentiality analysis results by both methods were combined. Finally, we demanded that for a gene to considered essential for growth on LB agar or LB broth the following three criteria should be met: (i) the gene is essential in LB-III by Tn-Seq Explorer analysis (ii) the gene is essential in LB-III by TRANSIT analysis (iii) the gene is essential in at least 5 out of the 6 analysis that was performed for the LB-I, LB-II, and LB-III by the two analysis tools (Additional file 1). We made an exception for 17 genes to be considered essential: requirements for essential genes in at least 5 out of the 6 analysis were reduced to 4. This exception was based on the other analysis for the same libraries selected and analyzed under different growth conditions. The same strategy was used to identify essential genes under Dip250-I, Dip250-II, and Dip400.

### Calculation of fitness for selected essential genes

Fitness of essential genes for all iron-restricted conditions were analyzed by using TRANSIT, with resampling option. LB-I was used as the input for Dip100 and Dip150, while LB-II was used as the input for Dip250-I, Dip250-II, and Dip400. For data normalization for fitness calculation, Trimmed Total Reads (TTR) option in TRANSIT was used and 10,000 samples were used for the analysis. The 5% of N-terminal and 10% of C-terminal of ORFs were removed and the gene fitness was considered if *p* value was ≤ 0.05.

## Acknowledgments

This research was supported by the Arkansas High Performance Computing Center (AHPCC) which is funded through multiple National Science Foundation grants and the Arkansas Economic Development Commission.

## Availability of data and materials

All Tn-seq sequencing data are available on NCBI Sequence Read Archive under BioProject number PRJNA397775.

## Authors’ contributions

Conceived and designed the experiments: YK SK. Performed the experiments, analyzed the data, wrote the manuscript: SK. Revised the manuscript: YK SK.

## Additional files

**Additional files 1: Algorithms utilized for essential gene identification.** Two tools were used for essential gene analysis, TRANSIT (Gumbel) and Tn-Seq Explorer. LB-I, LB-II, and Lb-III were analyzed separately by both tools for identification of essential genes. The gene was considered essential if 5 out of 6 analyses was essential (E) or essentiality index (EI) < 3. LB-I was transposon library inoculum subjected to Tn-seq without growth. LB-II and LB-III were grown in LB broth till mid-log phase. Dip is abbreviation of iron chelator 2,2’-Dipyridyl in μM. The number with Dip is concentration of Dip. (PDF)

Additional files 2: Full list of *Salmonella* Typhimurium essential genes in iron-replete (LB) conditions identified by Tn-seq. **(XLSL)**

**Additional files 3: The overlapped essential genes that identified by our Tn-seq method of *Salmonella* Typhimurium 14028S in iron-replete (LB) conditions and essential genes that identified by TraDIS method in *S*. Typhimurium SL3261.** (XLSL)

**Additional files 4: Required essential pathways of *Salmonella* Typhimurium in rich media.** Tn-seq libraries were grown in LB broth media till mid-log phase and on LB agar. The total number of essential genes were 336 and KEGG pathway analysis categorized them to 23 essential pathways. (PDF)

**Additional files 5: Full list of *Salmonella* Typhimurium essential genes in iron-restricted conditions identified by Tn-seq (Fe-independent essential genes).** Iron chelator 250 and 400 μM 2,2’-Dipyridyl were used. (XLSL)

**Additional files 6: Full list of *Salmonella* Typhimurium essential genes in iron-restricted conditions identified by Tn-seq (Fe-dependent essential genes).** Iron chelator 250 and 400 μM 2,2’-Dipyridyl were used. (XLSL)

Additional files 7: Average sequencing read counts in essential genes of *Salmonella* Typhimurium in iron-replete (LB) and iron-restricted (Dip400) conditions identified by Tn-seq. **(XLSL)**

**Additional files 8: Iron-dependent essential genes are essential genes in other bacteria.**

Almost all orthologous iron-dependent essential genes are genuine essential genes in (Keio) *E. coli*, *Salmonella Typhimurium* SL326, *Bacillus subtilis*, *Mycoplasma mycoid* JCVI-syn3.0, *Mycobacterium tuberculosis* H37Rv, and (LB-III) *Salmonella Typhimurium* 14028S in stress free conditions. (XLSL)

**Additional files 9: Bias in read sequencing is in favor of LB (iron-replete) conditions.** Total sequencing reads and insertions in ORFs are shown. Sequencing reads consumed by two mutants, *umuC* and *STM14_2428* are also shown. An insertion in iron-replete (LB) has a chance to get 227 sequence reads but this reduced to 100 reads in iron-restricted condition (Dip400). (XLSL)

