## Supplementary Materials for "Iron-dependent essential genes in *Salmonella* Typhimurium"

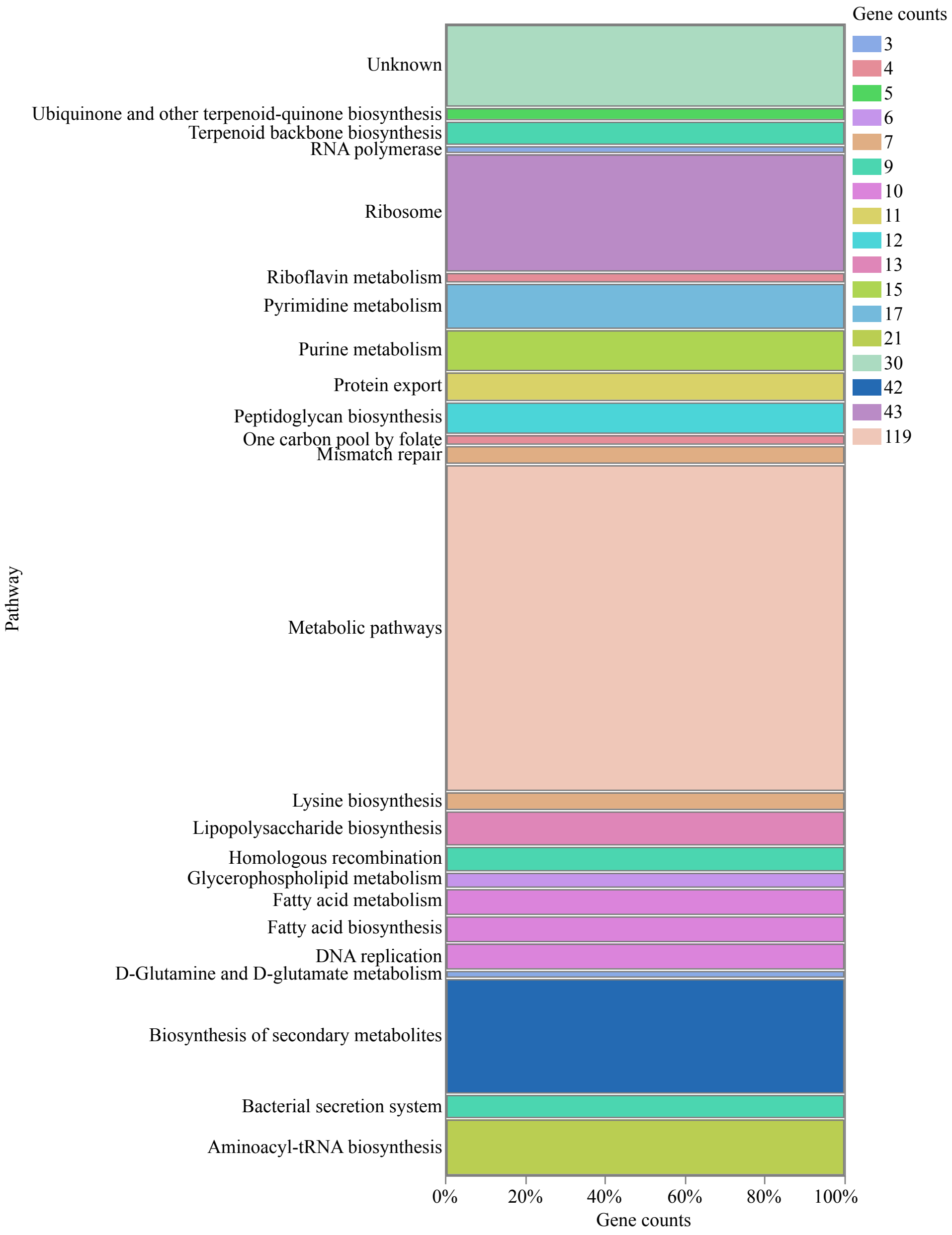


**Additional file 4. Required essential pathways of *Salmonella* Typhimurium in rich media.** Tn-seq libraries were grown in LB broth media till mid-log phase and on LB agar. The total number of essential genes were 336 and KEGG pathway analysis categorized them to 23 essential pathways.
