## Supplementary Materials for "Iron-dependent essential genes in *Salmonella* Typhimurium"

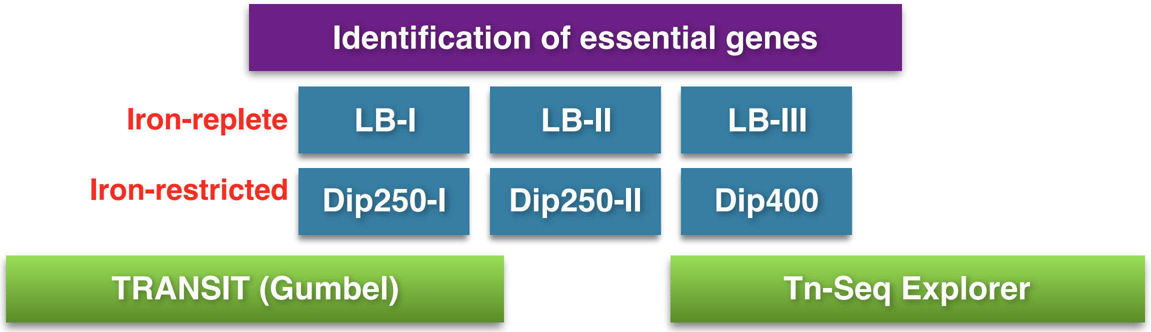

| | | | | |
|---|---|---|---|---|
| | | | A | LB-III Essentiality Index < 3 |
| B | LB-III Essential p < 0.05 | | | |
| | | C | | |
| | The gene has to be essential in at least 5 of the six essentiality analyses | | | |

Example-1

| Gene | LB-I | LB-II | LB-III | | LB-I | LB-II | LB-III | Call |
|---|---|---|---|---|---|---|---|---|
| *murA* | E | E | E | | 0 | 0 | 0 | E |
| *dapD* | NE | E | E | | 1 | 1 | 0 | E |
| *acpP* | NE | NE | E | | 2 | 1 | 1 | NE |
| *gmhA* | NE | NE | E | | 5 | 1 | 0 | NE |
| *ssb* | E | NE | E | | 3 | 0 | 1 | NE |
| *cydC* | E | E | E | | 3 | 0 | 1 | E |

| Gene | Dip250-I | Dip250-II | Dip400 | | Dip250-I | Dip250-II | Dip400 | Call |
|---|---|---|---|---|---|---|---|---|
| *murA* | NE | E | E | | 4 | 2 | 4 | NE |
| *dapD* | E | NE | E | | 0 | 2 | 2 | E |
| *acpP* | NE | E | NE | | 2 | 1 | 2 | NE |
| *gmhA* | NE | E | NE | | 1 | 0 | 1 | NE |
| *ssb* | NE | NE | E | | 4 | 2 | 3 | NE |
| *cydC* | E | E | E | | 2 | 4 | 5 | NE |

**Additional file 1. Algorithms utilized for essential gene identification.** Two tools were used for essential gene analysis, TRANSIT (Gumbel) and Tn-Seq Explorer. LB-I, LB-II, and Lb-III were analyzed separately by both tools for identification of essential genes. The gene was considered essential if 5 out of 6 analyses was essential (E) or essentiality index (EI) < 3. LB-I was transposon library inoculum subjected to Tn-seq without growth. LB-II and LB-III were grown in LB broth till mid-log phase. Dip is abbreviation of iron chelator 2,2`-Dipyridyl in µM. The number with Dip is concentration of Dip.
